# Autophagy modulation enhances enzyme replacement therapy response in Fabry disease

**DOI:** 10.1101/2021.04.26.441451

**Authors:** Lénaig Abily-Donval, Frédéric Barbey, Stéphanie Torre, Céline Lesueur, Marc Chevrier, Tony Pereira, Carine Pilon, Stéphane Marret, Abdellah Tebani, Soumeya Bekri

## Abstract

**Summary:** Fabry disease is a lysosomal disease due to α-galactosidase A (a-GalA) deficiency. Since 2001, Enzyme replacement therapy (ERT) has been used as specific treatment of Fabry disease, with variable effects depending on patient gender and affected organs. In Fabry cells, the endososomal/lysosomal system is highly altered. Consequently, the exogenous enzyme may be mistargeted and trapped into intracellular vesicles instead of reaching the lysosomes. In this study, we aimed to investigate the mechanisms underlying the processing of the exogenous enzyme. We used Fabry cells (cultured fibroblasts and podocyte cell line) to study the enzyme internalization and its effects on the catabolism of the main a-Gal A substrate, globotriaosylceramide (Gb3), upon autophagy inhibition. The exogenous enzyme reaches the early endosome in a similar timeframe in Fabry and control cells, while its targeting to lysosomes is delayed in Fabry cells. Gb3 concentration is lowered upon therapeutic enzyme addition or wortmannin treatment with a synergetic effect. These findings illustrate the positive impact of autophagy inhibition on enzyme trafficking and processing, allowing the increase of functional enzyme rate within the lysosome. Given the high cost of ERT, a better understanding of the cellular fate of the exogenous enzyme may lead to improve its targeting to the lysosome.

## 1. Introduction

Lysosomal diseases (LDs) are due to a lysosomal protein deficiency which leads to the accumulation of undegraded substrates within the lysosome. A combination of pathogenic mechanisms underlying LDs may interplay and disrupt the cellular network homeostasis. Lysosomes have a major role in the clearance of numerous endogenous and exogenous macromolecules [1,2]. Enlarged lysosomes may fail to play their proper role in major metabolic pathways such as the coordinated control of lysosome biogenesis [3], the orchestration of endosomal/lysosomal network [4] or the execution of autophagy process [2,5]. Several therapeutic strategies have been developed for LDs these last decades such as enzyme replacement (ERT), substrate reduction or gene therapies [2]. ERT has been applied for several LDs [6-8]. The concept relies on the uptake of exogenous enzymes through mannose-6-receptor-mediated endocytosis. Despite some successful results of the ERT, clinical efficacy has been challenged depending to the targeted tissue and the treated LD. These limitations were partly attributed to adverse immune responses and the production of antibodies against the exogenous enzyme that can account for efficacy reduction [3]. Another major drawback is the limited enzyme access to some potentially altered organs such as the central nervous system and the bones. Besides, even in the reachable organs, enzyme targeting to the lysosome may vary. These variations may be related to the trafficking and the processing of the recombinant enzyme from the cell membrane to the lysosome. Considering that the exogenous enzyme is directed into the endosome/lysosome system through a mannose-6-phosphate-mediated endocytosis and that this system is altered in LDs, enzyme targeting may be hampered and a significant proportion of the endocytosed enzyme may not reach the lysosomes. Indeed, it has been demonstrated that the autophagic block observed in LDs with the subsequent autophagic vacuole accumulation within the cells may prevent the enzyme from reaching the lysosomes [9]. If the autophagic buildup is prominent, the efficacy of exogenous enzyme may be significantly reduced [10,11]. Autophagy is a pivotal biological process in maintaining cellular homeostasis [12]. Lysosomes are involved in the different autophagic pathways namely, macroautophagy, chaperone-mediated autophagy and microautophagy. The major and most studied autophagic pathway, macroautophagy (named autophagy), is altered in LDs [5,10,13,14]. This pathway involves the biogenesis and fusion of several vesicles. The cellular components to be degraded are sequestered into a vesicle named autophagosome which fuses with a late endosome or a lysosome to form an amphisome ou an autolysosome respectively. The autophagosome content is thus digested by the lysosomal enzymes. Several key steps are required to the proper execution of this process : (i) the autophagosome initiation, (ii) lysosome biogenesis and integrity, (iii) autophagosomal fusion (iv) lysosomes reformation [15]. The autophagosome initiation involves a large set of autophagy related genes (ATG) and is tightly regulated through two main pathways, mTOR (mechanistic target of rapamycin) and Beclin1. Transcription factor EB (TFEB), plays a major role in lysosomal homeostasis by regulating the expression of numerous genes involved in lysosomes biogenesis and autophagy [16,17]. Autophagosome maturation and the endosomal/lysosomal vesicle fusion require large and coordinated machinery including ATG proteins and Rab GTPases [18].

Fabry disease is an X-linked LD due to a complete or partial α-galactosidase A deficiency, corresponding to the classic or multisystemic, severe form of the disease, or to the late-onset form. This disease is characterized by progressive accumulation of glycosphingolipids, mainly globotriaosylceramide (Gb3), in cells of many tissues [2]. First clinical manifestations appear during the childhood (neuropathy, skin lesions, cornea verticillata) then, in adulthood, multivisceral complications occur gradually (hypertrophic cardiomyopathy, renal failure and recurrent strokes) [19].

Since 2001, ERT is the gold standard treatment of Fabry disease. Two therapeutic enzymes are available: agalsidase alpha (Replagal®, Takeda); and agalsidase beta (Fabrazyme®, Sanofi). Although significant clinical benefits and a reduction of Gb3 levels in different biofluids and tissues have been reported in patients treated with ERT, the effects are variable depending on the patient gender and age, and on the affected organs [20]. A better understanding of the determinants of an efficient targeting of exogenous enzyme may allow circumventing variable ERT efficacy.

In this study, we aimed to investigate the internalization of therapeutic enzyme and its targeting within the lysosome in Fabry cells and the effect of autophagy modulation on ERT efficacy.

## 2. Materials and Methods

### 2.1. Cell culture, treatment and analysis

#### 2.1.1. Fibroblasts

A skin biopsy was performed in five patients with the classic form of Fabry disease, followed in the Centre Hospitalier Universitaire Vaudois (CHUV), in Lausanne. Each of the five patients and controls included in this study provided written and signed informed consent. The study was performed according to the Declaration of Helsinki. These patients were treated with Agalsidase alpha 0,2 mg/Kg intravenously every 2 weeks. Control fibroblasts were obtained from five healthy male individuals. Fibroblasts were maintained in 80% of Ham’s medium (Biochrom AG) supplemented with 10% of fetal bovine serum –FBS-(Eurobio), 1% of penicillin 100 UI/ml and streptomycin 100µg/ml (Sigma-Aldrich) and 1% of glutamine, and 20% of Chang medium (IrvineScientific) supplemented with 1% of penicillin 100 UI/ml and streptomycin 100µg/ml. All cells were grown at 37°C with 5% CO_2_. Fibroblasts are cultured on 100 millimeters diameter dishes in (i) standard conditions (ii) with or without recombinant enzyme Agalsidase alpha (5µl/ml) (Takeda) and (iii) with or without wortmannin (75nM) (Sigma-Aldrich). Of note, the cells have been pre-treated with wortmannin 24 hours before enzyme addition. For immunohistology studies, fibroblasts were cultured on 12 millimeters diameter dishes, with or without 2,5 µl/ml of fluorescent enzyme (eGFP-α galactosidase A) at 24 and 72 hours, with or without wortmaninn.

#### 2.1.2. Podocytes

Immortalized human podocyte cell line generated using CRISPR/Cas9 technology was provided by Ora Weisz Laboratory, University of Pittsburgh School of Medicine. This cell line is a validated model for Fabry disease [21]. The wild-type podocytes were used as control cell line. These cells were cultured in DMEM/F12 medium (Gibco®) supplemented with 10% of fetal bovine serum –FBS-(Eurobio), 1% of penicillin 100 UI/ml and streptomycin 100µg/ml (Sigma-Aldrich) and 1% of glutamine. All cells were grown at 37°C with 5% CO_2_. Podocytes are cultured on 100 millimeters diameter dishes in (i) standard conditions (ii) with or without Agalsidase alpha (5µl/ml) (Takeda) and (iii) with or without wortmannin (75nM) (Sigma-Aldrich). As for the fibroblasts, the cells have been pre-treated with wortmannin 24 hours before enzyme addition.

### 2.2. Western blot analysis

Different endosomal/lysosomal vesicles are involved in the trafficking and processing of exogenous enzyme and its targeting to the lysosomes. Early endosomal autoantigen 1 (EEA1) is an early endosome biomarker involved as a membrane tethering factor in the fusion of early endososomes during the endocytosis [22]. Rab GTPase proteins play a pivotal role in the regulation of cellular trafficking. These proteins localize to the specific organelle and regulate and thus interact directly with their effectors. Rab9 is linked to the late endososome and mediates the cation-independent mannose-6-phosphate receptor recycling [23]. Rab11 regulates the recycling of endocytosed proteins [24]. Rab7 controls the maturation of early endosome to late endosomes and is required for the autophagosome-lysosome fusion and lysosome positioning within the cell [24-26]. We, therefore, quantified EEA1, Rab11, Rab9 and Rab7 by western blot analysis with the indicated antibodies.

Fibroblasts and podocytes were washed twice with PBS and proteins were extracted with the lysis buffer (Cell Signaling Technology) which protease and phosphatase inhibitors (Sigma-Aldrich). To analyze protein levels, the protein pellets were solubilized in Laemmli buffer. Fifty μg of proteins from fibroblast samples were analyzed by SDS-PAGE 10% under reducing conditions and transferred to nitrocellulose membrane. The membrane was then blocked with 1X TBS, 0.5% tween-20, 5% nonfat milk and incubated with EEA1, Rab11, Rab9 and Rab7 antibodies (Abcam). Enhanced chemiluminescence reagent ECL (GE Healthcare Limited) was used for protein detection with HRP-conjugated anti-mouse antibody (Santa Cruz). For normalization, membrane was dehybridized by incubating the membrane 30 min at 50°C in a dehybridization solution (Tris-HCL 0.5 M pH 6.8, SDS 10%, mercaptoethanol 7 mM) and incubated with mouse monoclonal anti-β-actin (Sigma-Aldrich). Quantification of the images was performed using Quantity one and Gel doc XR from Biorad (Hercules).

### 2.2. Immunochemistry studies

Rabbit polyclonal anti-EEA1 antibody (LifeSpan Biosciences) was used at a dilution of 1/100 in a solution of PBS -0.03 % triton and BSA 1 %. Secondary antibodies conjugated to Alexa 594 were used at 1/300 in a solution of PBS -0.03 % triton and BSA 1%. Acidophilic LysoTrackerTM Red stain (Invitrogen) was added to the cell culture in culture medium after 72h of treatment and incubated for 2 hours. The immunofluorescence images were obtained using the Leica laser-scanning confocal image system SP8 (Leica Microsystems AG).

### 2.3. Gb3 assessment using mass spectrometry

Gb3 measurement was implemented according to Mills’ method [27]. The primary step is a liquid phase extraction from cell residues. Cells were scraped with phosphate-buffered saline (PBS) and centrifuged 10 minutes at 2000 revolutions per minute. The pellet was hydrated using 500 µl of water (Water Plus, Carlo Erba) and mixed using ultrasonic sound 5 times for 2 seconds. Three hundred µL of this mix were added to 300 µl of water (Water Plus, Carlo Erba) and 50 µl of C17-CTH (Matreya) at 0.05 µg/ml used as calibrator. A sample of 100µl was used for protein assessment. Extraction was realized with the addition of 5 mL of dichloromethane (Merck): methanol (Carlo Erba) (2:1, v/v) by shaking on multivortexer for 10 minutes. The two layers were separated by centrifugation for 5 minutes at 3500 revolutions per minute. The higher layer was transferred to a new vial and dried under N_2_ at 40°C. Prior to mass spectrometer analysis, 100µl of methanol were added. Gb3 quantitative analysis was performed with the mass spectrometer Sciex 4000 QTRAP (Sciex) using the electrospray ion source (TurbolonSpray). Ten microliters were directly injected into the electrospray via HPLC line (Shimadzu) in FIA mode (flow injection analysis).

### 2.4. Statistical analysis

Data are described as mean with standard deviation. A one-way analysis of variance (ANOVA) test was applied for multiple groups testing while a t-test is used for binary comparisons. The Benjamini and Hochberg false discovery rate (FDR) method was used for multiple testing corrections with an FDR cut-off level of 5%. All statistical analyses and visualization were done with the R software.

## 3. Results

### 3.1. Endosomal/lysosomal biomarker expression in Fabry cells

To evaluate the endocytic alteration in Fabry cells that may hamper the lysosomal targeting of exogenous enzyme, we studied the intracellular endosomal/lysosomal biomarker expression in control and Fabry fibroblasts.

No significant changes in EEA1, Rab11 and Rab9 levels have been observed between control and Fabry cells, while Rab7 amount was significantly higher in Fabry cells compared to control cells (Figure 1). These results are in line with the autophagosomal maturation defect observed in Fabry disease [28].

**Figure 1.**
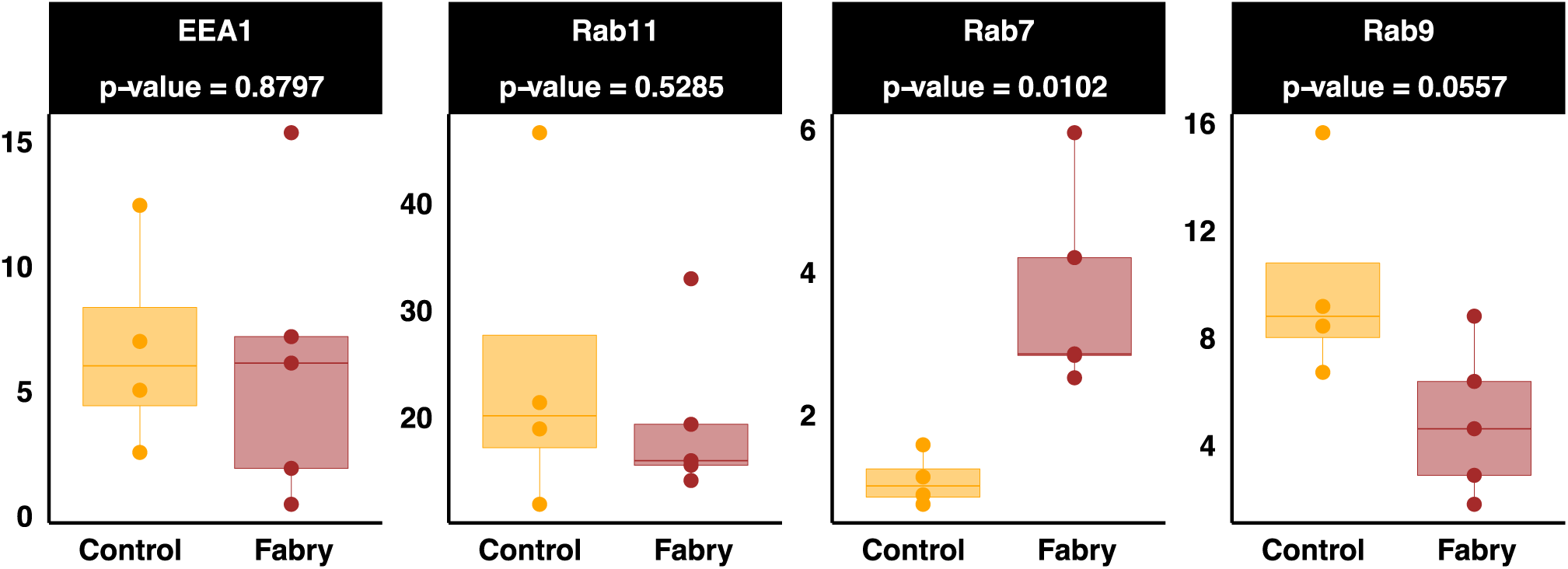
Endosomal/lysosomal biomarker expression in control and Fabry fibroblasts. Quantitative analysis of autophagic activity by Western blot technique. Fifty µg of protein from fibroblasts were analyzed by SDS-PAGE 10% under reducing conditions. EAA1, Rab9, Rab11 and Rab7 were detected by immunoblotting. β-actin was used as an internal control. Statistical significance is set at p > 0.05.

### 3.2. Exogenous enzyme trafficking and processing

The internalization course of eGFP-α galactosidase A 12 and 72h after enzyme addition and its co-localization with EEA1, an early endosome biomarker, provide similar results in control and Fabry cells (Figure 2A). Indeed, the eGFP-α galactosidase A and EEA1 signals showed an extensive co-localization 12 h after enzyme treatment, while there is virtually no fluorescence merged signals after 72 hours. Thus, exogenous enzyme internalization to the early endosome in Fabry cells is not altered.

**Figure 2.**
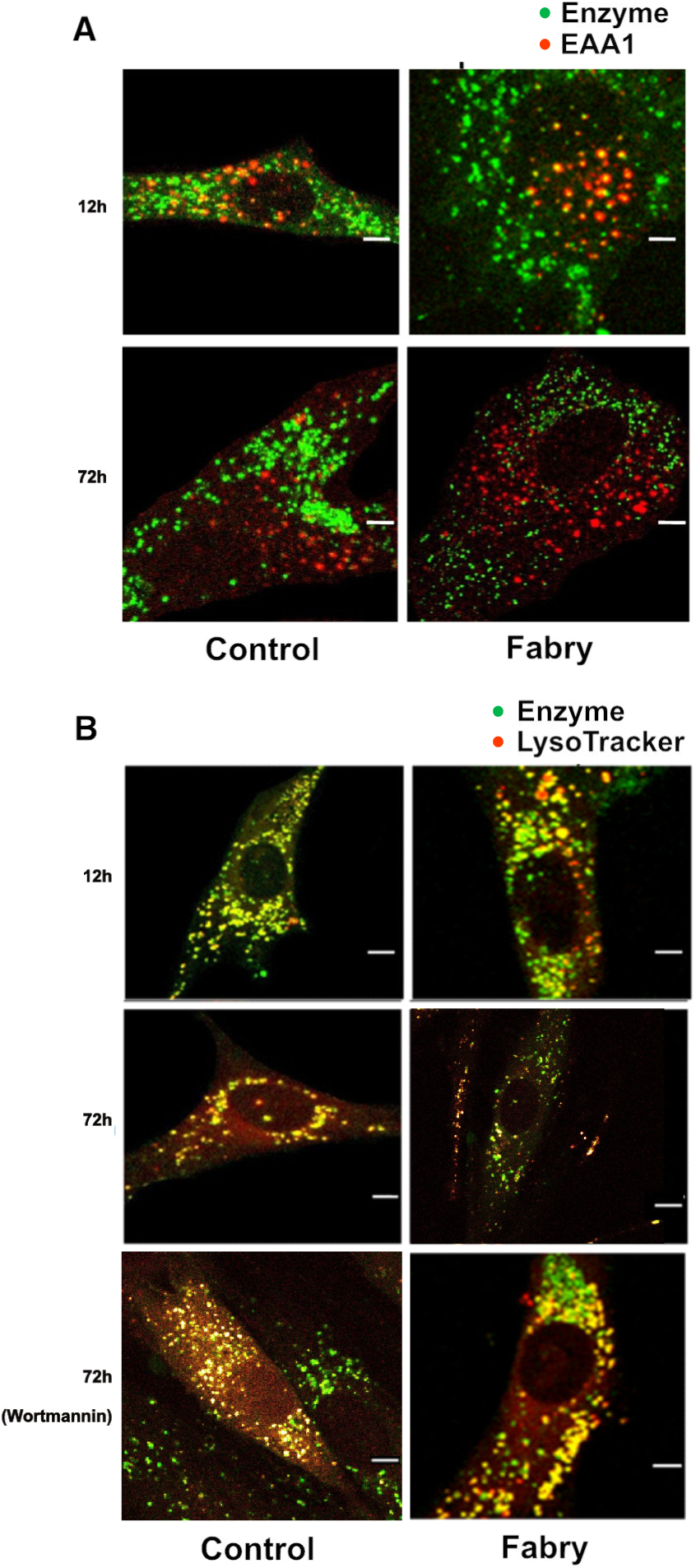
Internalization of the recombinant enzyme in the early endosome and its targeting within the lysosome. A. Confocal microscopy of control and Fabry fibroblasts after eGFP-α galactosidase A (green -2,5µL/mL) incubation for 12 or 72 h. Cultured cells were immunolabelled with anti-EEA1 antibody (red). The immunofluorescent images were obtained using a laser-scanning confocal image system (Scale bar = 10 µM). B. Confocal microscopy of control and Fabry cells after eGFP-α galactosidase A incubation (incubation time: 12 or 72 h). Cultured cells were incubated with LysotrackerTM. The immuno-fluorescent images were obtained using a laser-scanning confocal image system (Scale bar = 10µM).

### 3.3. Lysosomal targeting of the exogenous enzyme is delayed in Fabry cells

In control cells, eGFP-α galactosidase A and LysotrackerTM signals co-localized 72h following enzyme incubation. In contrast green staining corresponding to eGFP-α galactosidase A is still present in Fabry cells 72 h after enzyme treatment. These results are consistent with a delayed targeting of the exogenous enzyme to the lysosomes in Fabry cells compared to control cells (Figure 2B). Intracellular trafficking seems to be altered in Fabry disease which may explain the low rate of exogenous enzyme reaching the lysosomes. Wortmannin addition increased the enzyme and lysosome colocalization (Figure 2B).

### 3.4. Autophagy modulation enhances the Gb3 cleavage in Fabry cells incubated with exogenous enzyme

The activity of exogenous a-Gal A efficiently internalized and processed in Fabry cells was evaluated using the cellular Gb3 assessment. Control and Fabry cells were subjected or not to autophagy inhibition through wortmannin treatment. In control cells, no difference in Gb3 concentration has been observed between standard condition, enzyme treatment, wortmannin and wortmannin/enzyme conditions. As expected in Fabry cells, a significantly higher amounts of Gb3 were present in basal condition compared to control cells (Figure 3A), enzyme as well as wortmannin addition succeeded in reducing this amount significantly in both fibroblasts and podocytes, this reduction was heightened in presence of both enzyme and wortmannin indicating a synergic effect of these two molecules on ERT trafficking and processing (Figure 3A, Table 1). The visualization of the results for each fibroblast sample as line plots illustrates the change of Gb3 concentration patterns in cell samples subjected to different conditions and showed that this synergetic effect is found for all studied samples (Figure 3B).

**Figure 3.**
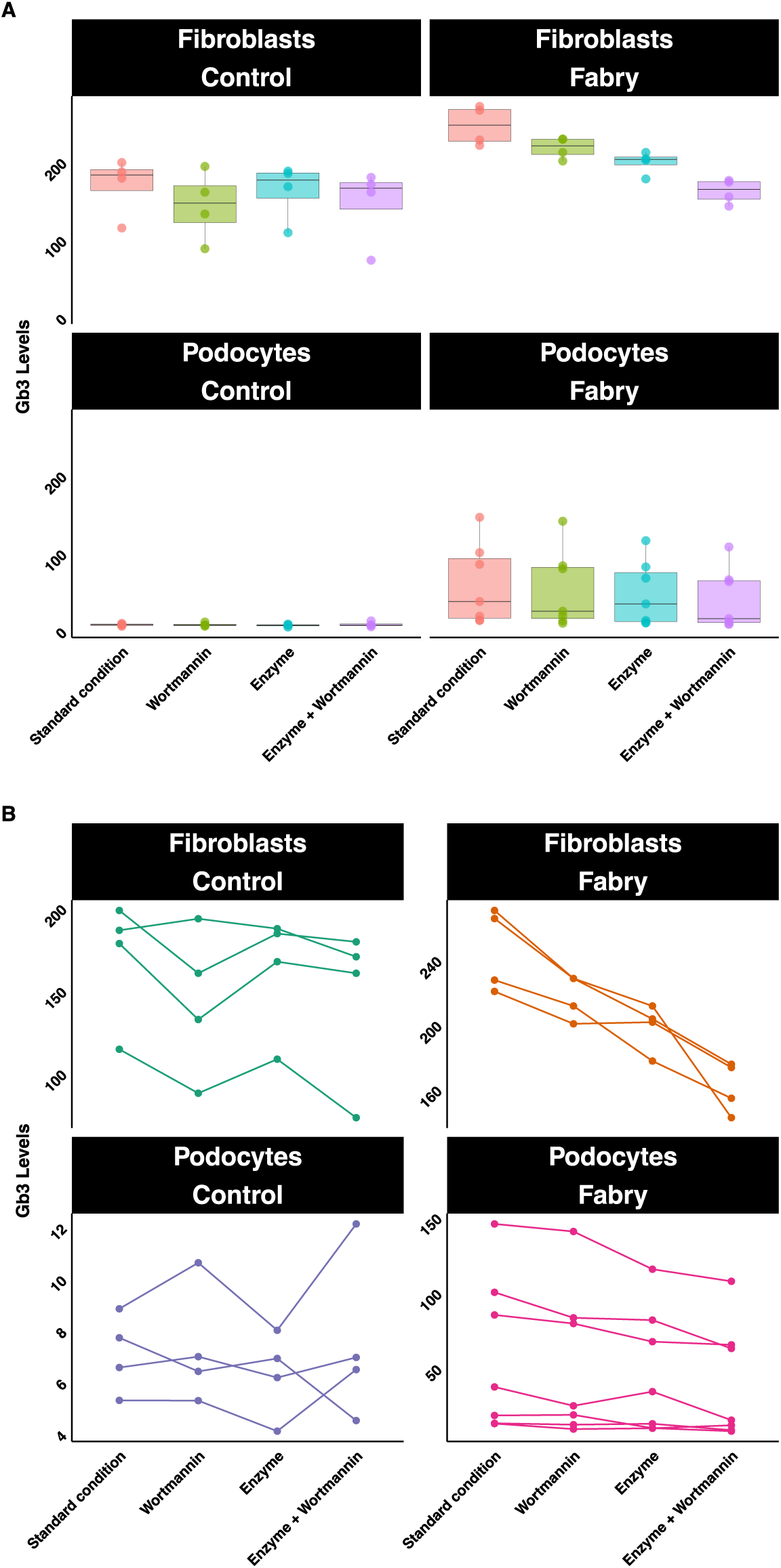
Effect of autophagy modulation on Gb3 cleavage in control and Fabry cells. Cellular Gb3 assessment using control and Fabry fibroblasts or Fabry podocytes subjected or not to autophagy inhibition through wortmannin treatment. A. Boxplots are used to visualize the spread of the Gb3 concentrations across the different conditions (basal, enzyme, wortmannin, enzyme/wortmannin). B. Line plots are used to visualize the change of Gb3 concentration patterns in samples subjected to different conditions (basal, wortmannin, enzyme, enzyme/wortmannin). Detailed statistical metrics are presented in Table 1.

**Table 1.**
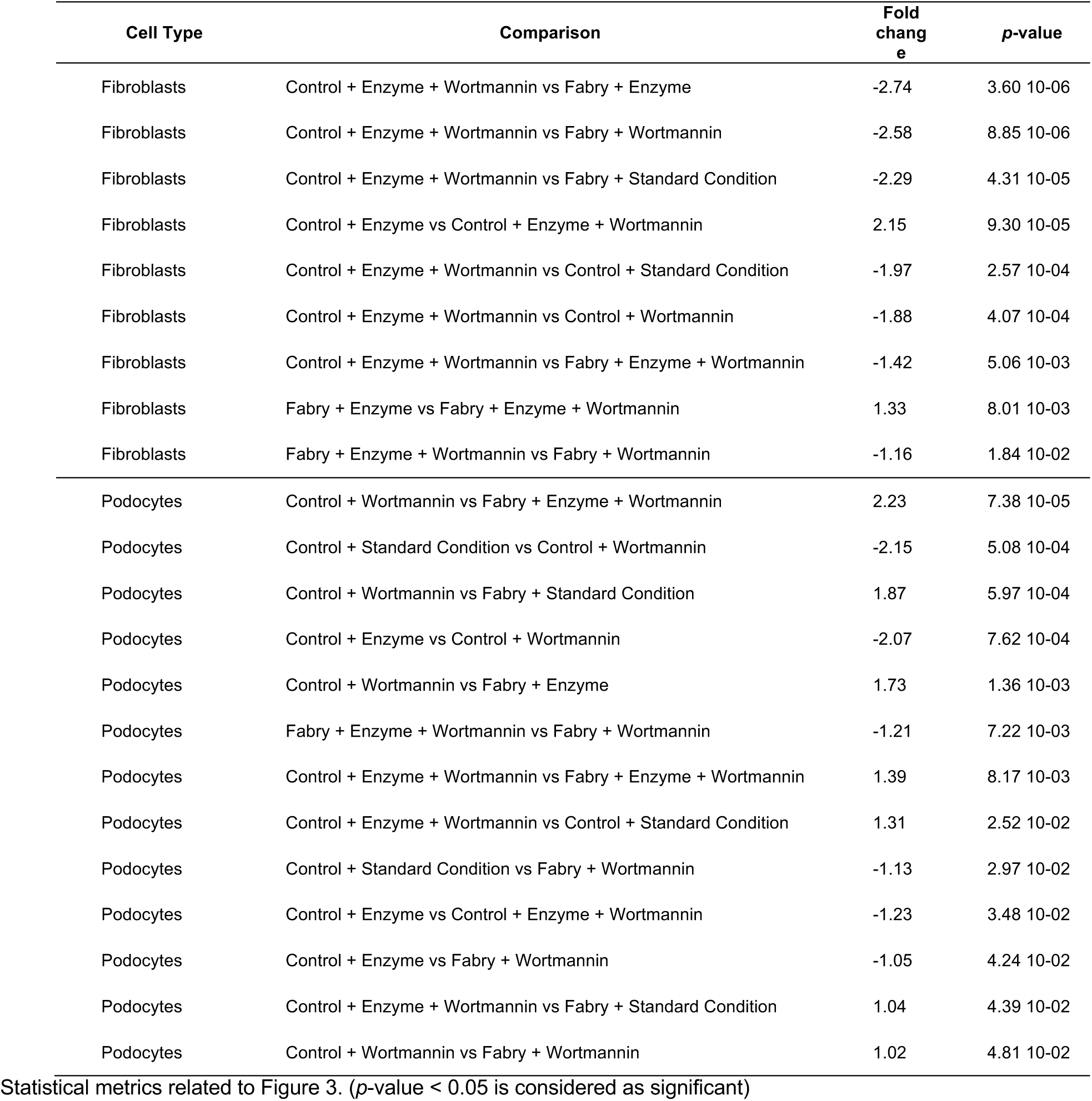
Effect of autophagy modulation on Gb3 cleavage in control and Fabry cells.

| Cell Type | Comparison | Fold change | p-value |
| --- | --- | --- | --- |
| Fibroblasts | Control + Enzyme + Wortmannin vs Fabry + Enzyme | -2.74 | 3.60 10<sup>-06</sup> |
| Fibroblasts | Control + Enzyme + Wortmannin vs Fabry + Wortmannin | -2.58 | 8.85 10<sup>-06</sup> |
| Fibroblasts | Control + Enzyme + Wortmannin vs Fabry + Standard Condition | -2.29 | 4.31 10<sup>-05</sup> |
| Fibroblasts | Control + Enzyme vs Control + Enzyme + Wortmannin | 2.15 | 9.30 10<sup>-05</sup> |
| Fibroblasts | Control + Enzyme + Wortmannin vs Control + Standard Condition | -1.97 | 2.57 10<sup>-04</sup> |
| Fibroblasts | Control + Enzyme + Wortmannin vs Control + Wortmannin | -1.88 | 4.07 10<sup>-04</sup> |
| Fibroblasts | Control + Enzyme + Wortmannin vs Fabry + Enzyme + Wortmannin | -1.42 | 5.06 10<sup>-03</sup> |
| Fibroblasts | Fabry + Enzyme vs Fabry + Enzyme + Wortmannin | 1.33 | 8.01 10<sup>-03</sup> |
| Fibroblasts | Fabry + Enzyme + Wortmannin vs Fabry + Wortmannin | -1.16 | 1.84 10<sup>-02</sup> |
| Podocytes | Control + Wortmannin vs Fabry + Enzyme + Wortmannin | 2.23 | 7.38 10<sup>-05</sup> |
| Podocytes | Control + Standard Condition vs Control + Wortmannin | -2.15 | 5.08 10<sup>-04</sup> |
| Podocytes | Control + Wortmannin vs Fabry + Standard Condition | 1.87 | 5.97 10<sup>-04</sup> |
| Podocytes | Control + Enzyme vs Control + Wortmannin | -2.07 | 7.62 10<sup>-04</sup> |
| Podocytes | Control + Wortmannin vs Fabry + Enzyme | 1.73 | 1.36 10<sup>-03</sup> |
| Podocytes | Fabry + Enzyme + Wortmannin vs Fabry + Wortmannin | -1.21 | 7.22 10<sup>-03</sup> |
| Podocytes | Control + Enzyme + Wortmannin vs Fabry + Enzyme + Wortmannin | 1.39 | 8.17 10<sup>-03</sup> |
| Podocytes | Control + Enzyme + Wortmannin vs Control + Standard Condition | 1.31 | 2.52 10<sup>-02</sup> |
| Podocytes | Control + Standard Condition vs Fabry + Wortmannin | -1.13 | 2.97 10<sup>-02</sup> |
| Podocytes | Control + Enzyme vs Control + Enzyme + Wortmannin | -1.23 | 3.48 10<sup>-02</sup> |
| Podocytes | Control + Enzyme vs Fabry + Wortmannin | -1.05 | 4.24 10<sup>-02</sup> |
| Podocytes | Control + Enzyme + Wortmannin vs Fabry + Standard Condition | 1.04 | 4.39 10<sup>-02</sup> |
| Podocytes | Control + Wortmannin vs Fabry + Wortmannin | 1.02 | 4.81 10<sup>-02</sup> |

Statistical metrics related to Figure 3. ( $p$ -value < 0.05 is considered as significant)

## 4. Discussion

The concept of enzyme replacement therapy as specific treatment of LDs is appealing. It consists in bringing a functional exogenous enzyme into the lysosome to fix the primary deficit. However, the trafficking and processing of this enzyme to the lysosome rely on endosomal/lysosomal system. Hypothetically, if the endososomal/lysosomal system is highly altered and an autophagic build-up is constituted, the therapeutic enzyme may be mistargeted and trapped into intracellular vesicles instead of reaching the lysosomes. Given the high cost of ERT, a better understanding of the cellular fate of the exogenous enzyme may lead to improve its targeting to the lysosome, the unique organelle where it could exert its biological function. Identifying novel therapeutic interventions that could enhance ERT efficacy may help improving the clinical features and lowering the costs.

In this study, we used cultured fibroblasts from Fabry patients and control individuals, and podocyte cell line invalidated for *GLA* gene and control podocyte cell line to study exogenous enzyme trafficking and processing. Protein expression of different endosomal biomarkers was studied in Fabry versus control cells. Rab7 level was significantly elevated in Fabry fibroblasts compared to control. Rab7 controls the autophagosome/lysosomes fusion and is involved in lysosome positioning [26]. It has been demonstrated that the peripheral lysosomes within the cell have a higher intraluminal pH than the lysosome located in the perinuclear area and the lysosomes mobility relies on Rab7 [26]. In addition, Wong et al. reported recently that the contact sites between lysosomes and mitochondria depend on Rab7 [29]. Considering the multiple effects of Rab7, it appears that the changes in Rab7 concentration may be challenging for cell homeostasis. This elevated concentration may disrupt the above-mentioned mechanism.

The first step of exogenous enzyme endocytosis seems to be preserved in Fabry cells. The enzyme is located within the early endosome 12 h after the enzyme addition, while after 72 h, the enzyme staining is distinct from that of the early endosome, illustrating that the enzyme has moved from the early endosome to other compartments. Importantly, our results showed that the lysosomal targeting of the exogenous enzyme is delayed in Fabry patients. After 72 h, the enzyme/lysosome signal co-localization rate is lower in Fabry cells compared to controls which indicates that a proportion of the exogenous enzyme did not reach the lysosomes. In order to confirm these results and to study the effect of autophagy modulation on ERT efficacy, a functional test has been performed by assessing the Gb3 cellular concentration 72 h after enzyme treatment. The addition of the exogenous and wortmannin, an autophagy inhibitor, succeeded in lowering Gb3 concentration. This finding illustrates the positive impact of autophagy inhibition on enzyme trafficking and processing allowing the increase of functional enzyme rate.

Moreover, the synergic effect of ERT and wortmannin is demonstrated by the significant reduction of cellular Gb3 concentration compared to the condition enzyme or wortmannin alone. The autophagy modulation in Fabry disease may, thus, act as adjunctive therapy and smoothen the ERT targeting.

Mistargeting the therapeutic enzyme has been described in Pompe disease and autophagy suppression by acting on mTor pathway resulted in a clearance of autophagy build-up and facilitates ERT targeting [30,31]. Indeed, the authors proposed a combination of mTor-mediated inhibition of autophagy such as arginine supplementation with ERT for the treatment of Pompe disease.

## 5. Conclusions

In conclusion, ERT might have the potential to address the enzyme deficiency consequences if its final destination is reached. Our experimental data indicate that in Fabry cells, therapeutic enzyme has a partial efficacy on Gb3 clearance and concomitant autophagy inhibition may represent an effective strategy for optimizing ERT treatment.

## Author Contributions

Conceptualization, S.B.; methodology, S.B., L.A.D.; formal analysis, L.A.D., S.T., C.L., C.P., T.P., A.T.; resources, F.B. and S.B.; writing—original draft preparation, S.B.; writing—review and editing, F.B., B.G., S.M. and A.T.; visualization, A.T.; supervision, S.B.; project administration, S.B.; funding acquisition, S.B. All authors have read and agreed to the published version of the manuscript.

## Institutional Review Board Statement

The study was conducted according to the guidelines of the Declaration of Helsinki and approved by the CER-VD, Lausanne, Switzerland (Commission cantonale d’éthique de la recherche sur l’être humain: protocole 101/01).

## Informed Consent Statement

Informed consent was obtained from all subjects involved in the study.

## Data Availability Statement

All the data that supports the findings are presented in the manuscript.

## References

1. Parenti, G., Medina, D.L., Ballabio, A. The rapidly evolving view of lysosomal storage diseases. EMBO molecular medicine 2021, 10.15252/emmm.202012836, e12836, doi:10.15252/emmm.202012836.

2. Platt, F.M., d’Azzo, A., Davidson, B.L., Neufeld, E.F., Tifft, C.J. Lysosomal storage diseases. Nature reviews. Disease primers 2018, 4, 27, doi:10.1038/s41572-018-0025-4.

3. Marques, A.R.A., Saftig, P. Lysosomal storage disorders - challenges, concepts and avenues for therapy: beyond rare diseases. Journal of cell science 2019, 132, doi:10.1242/jcs.221739.

4. Fukuda, T., Ewan, L., Bauer, M., Mattaliano, R.J., Zaal, K., Ralston, E., Plotz, P.H., Raben, N. Dysfunction of endocytic and autophagic pathways in a lysosomal storage disease. Annals of neurology 2006, 59, 700–708, doi:10.1002/ana.20807.

5. Segatori, L. Impairment of homeostasis in lysosomal storage disorders. IUBMB life 2014, 66, 472–477, doi:10.1002/iub.1288.

6. Li, M. Enzyme Replacement Therapy: A Review and Its Role in Treating Lysosomal Storage Diseases. Pediatric annals 2018, 47, e191–e197, doi:10.3928/19382359-20180424-01.

7. Safary, A., Akbarzadeh Khiavi, M., Mousavi, R., Barar, J., Rafi, M.A. Enzyme replacement therapies: what is the best option? BioImpacts : BI 2018, 8, 153–157, doi:10.15171/bi.2018.17.

8. Thomas, R., Kermode, A.R. Enzyme enhancement therapeutics for lysosomal storage diseases: Current status and perspective. Molecular genetics and metabolism 2019, 126, 83–97, doi:10.1016/j.ymgme.2018.11.011.

9. Lim, J.A., Li, L., Raben, N. Pompe disease: from pathophysiology to therapy and back again. Frontiers in aging neuroscience 2014, 6, 177, doi:10.3389/fnagi.2014.00177.

10. Fukuda, T., Ahearn, M., Roberts, A., Mattaliano, R.J., Zaal, K., Ralston, E., Plotz, P.H., Raben, N. Autophagy and mistargeting of therapeutic enzyme in skeletal muscle in Pompe disease. Molecular therapy : the journal of the American Society of Gene Therapy 2006, 14, 831–839, doi:10.1016/j.ymthe.2006.08.009.

11. Nascimbeni, A.C., Fanin, M., Tasca, E., Angelini, C., Sandri, M. Impaired autophagy affects acid alpha-glucosidase processing and enzyme replacement therapy efficacy in late-onset glycogen storage disease type II. Neuropathology and applied neurobiology 2015, 41, 672–675, doi:10.1111/nan.12214.

12. Jiang, P., Mizushima, N. Autophagy and human diseases. Cell research 2014, 24, 69–79, doi:10.1038/cr.2013.161.

13. Lieberman, A.P., Puertollano, R., Raben, N., Slaugenhaupt, S., Walkley, S.U., Ballabio, A. Autophagy in lysosomal storage disorders. Autophagy 2012, 8, 719–730, doi:10.4161/auto.19469.

14. Settembre, C., Fraldi, A., Jahreiss, L., Spampanato, C., Venturi, C., Medina, D., de Pablo, R., Tacchetti, C., Rubinsztein, D.C., Ballabio, A. A block of autophagy in lysosomal storage disorders. Human molecular genetics 2008, 17, 119–129, doi:10.1093/hmg/ddm289.

15. Fujioka, Y., Noda, N.N. Biomolecular condensates in autophagy regulation. Current opinion in cell biology 2021, 69, 23–29, doi:10.1016/j.ceb.2020.12.011.

16. Bajaj, L., Lotfi, P., Pal, R., Ronza, A.D., Sharma, J., Sardiello, M. Lysosome biogenesis in health and disease. Journal of neurochemistry 2019, 148, 573–589, doi:10.1111/jnc.14564.

17. Willett, R., Martina, J.A., Zewe, J.P., Wills, R., Hammond, G.R.V., Puertollano, R. TFEB regulates lysosomal positioning by modulating TMEM55B expression and JIP4 recruitment to lysosomes. Nature communications 2017, 8, 1580, doi:10.1038/s41467-017-01871-z.

18. Zhao, Y.G., Zhang, H. Autophagosome maturation: An epic journey from the ER to lysosomes. The Journal of cell biology 2019, 218, 757–770, doi:10.1083/jcb.201810099.

19. Tsujiuchi, M., Ebato, M., Maezawa, H., Mizukami, T., Nogi, A., Ikeda, N., Iso, Y., Suzuki, H. Long-Term Effects of Enzyme Replacement Therapy for Anderson-Fabry Disease. International heart journal 2019, 60, 208–214, doi:10.1536/ihj.17-688.

20. Azevedo, O., Gago, M.F., Miltenberger-Miltenyi, G., Sousa, N., Cunha, D. Fabry Disease Therapy: State-of-the-Art and Current Challenges. Int J Mol Sci 2020, 22, doi:10.3390/ijms22010206.

21. Oyarzun, J.E., Lagos, J., Vazquez, M.C., Valls, C., De la Fuente, C., Yuseff, M.I., Alvarez, A.R., Zanlungo, S. Lysosome motility and distribution: Relevance in health and disease. Biochimica et biophysica acta. Molecular basis of disease 2019, 1865, 1076–1087, doi:10.1016/j.bbadis.2019.03.009.

22. Ramanathan, H.N., Zhang, G., Ye, Y. Monoubiquitination of EEA1 regulates endosome fusion and trafficking. Cell & bioscience 2013, 3, 24, doi:10.1186/2045-3701-3-24.

23. Kucera, A., Borg Distefano, M., Berg-Larsen, A., Skjeldal, F., Repnik, U., Bakke, O., Progida, C. Spatiotemporal Resolution of Rab9 and CI-MPR Dynamics in the Endocytic Pathway. Traffic (Copenhagen, Denmark) 2016, 17, 211–229, doi:10.1111/tra.12357.

24. Welz, T., Wellbourne-Wood, J., Kerkhoff, E. Orchestration of cell surface proteins by Rab11. Trends in cell biology 2014, 24, 407–415, doi:10.1016/j.tcb.2014.02.004.

25. Bonifacino, J.S., Neefjes, J. Moving and positioning the endolysosomal system. Current opinion in cell biology 2017, 47, 1–8, doi:10.1016/j.ceb.2017.01.008.

26. Cabukusta, B., Neefjes, J. Mechanisms of lysosomal positioning and movement. Traffic (Copenhagen, Denmark) 2018, 19, 761–769, doi:10.1111/tra.12587.

27. Mills, K., Johnson, A., Winchester, B. Synthesis of novel internal standards for the quantitative determination of plasma ceramide trihexoside in Fabry disease by tandem mass spectrometry. FEBS letters 2002, 515, 171–176, doi:10.1016/s0014-5793(02)02491-2.

28. Chevrier, M., Brakch, N., Celine, L., Genty, D., Ramdani, Y., Moll, S., Djavaheri-Mergny, M., Brasse-Lagnel, C., Annie Laquerriere, A.L., Barbey, F., et al. Autophagosome maturation is impaired in Fabry disease. Autophagy 2010, 6, 589–599, doi:10.4161/auto.6.5.11943.

29. Wong, Y.C., Kim, S., Peng, W., Krainc, D. Regulation and Function of Mitochondria-Lysosome Membrane Contact Sites in Cellular Homeostasis. Trends in cell biology 2019, 29, 500–513, doi:10.1016/j.tcb.2019.02.004.

30. Lim, J.A., Li, L., Shirihai, O.S., Trudeau, K.M., Puertollano, R., Raben, N. Modulation of mTOR signaling as a strategy for the treatment of Pompe disease. EMBO molecular medicine 2017, 9, 353–370, doi:10.15252/emmm.201606547.

31. Lim, J.A., Sun, B., Puertollano, R., Raben, N. Therapeutic Benefit of Autophagy Modulation in Pompe Disease. Molecular therapy : the journal of the American Society of Gene Therapy 2018, 26, 1783–1796, doi:10.1016/j.ymthe.2018.04.025.

